# Brain gene expression in a novel mouse model of postpartum mood disorder

**DOI:** 10.1101/555870

**Authors:** Trevor Humby, William Davies

**Affiliations:** School of Psychology, Cardiff University, UK; Neuroscience and Mental Health Research Institute, Cardiff University, UK; Medical Research Council Centre for Neuropsychiatric Genetics and Genomics and Division of Psychological Medicine and Clinical Neurosciences, School of Medicine, Cardiff University, UK

**Keywords:** Female, mice, depression, postpartum, steryl-sulfatase, psychotic disorders, receptors, G-protein coupled

## Abstract

**Introduction:** Steroid sulfatase (STS) is an enzyme which cleaves sulfate groups from a variety of steroid hormones, thereby altering their activity and solubility. The expression and activity of STS is increased in female mammalian tissues (including brain) during late pregnancy and into the postpartum period. STS-deficient human and mouse mothers (as a consequence of genetic mutation or acute pharmacological manipulation) show evidence for elevated psychopathology and abnormal behaviour respectively in the postpartum period. In mice, these behavioural effects can be partially normalised through administration of the antipsychotic ziprasidone.

**Methods:** To explore the neurobiology underlying these postpartum behavioural effects, we compared whole brain gene expression by microarray in behaviourally-defined new mouse mothers acutely administered the STS inhibitor 667-Coumate (10mg/kg p.o.) or vehicle solution (n=12 per group); significant changes were followed-up with pathway analysis and quantitative polymerase chain reaction (qPCR). Finally, the effects of combined 667-Coumate and antipsychotic (ziprasidone) administration (0, 0.3 and 1.0mg/kg i.p.) on the brain expression of the most robustly differentially-expressed candidate genes was examined (n≥7 per group).

**Results:** Surprisingly, no significant gene expression changes were detected between vehicle and 667-Coumate-treated brains at a False Discovery Rate (FDR) corrected p<0.1. 1,081 unique expression changes were detected at a less-stringent cut-off of p<0.05, just two top hits were verified by qPCR, and pathway analysis indicated a significant enrichment of genes involved in olfactory transduction (corrected p-value=1.8×10^−3^). The expression of the two most robust differentially-expressed genes (*Stoml3* and *Cyp2g1*) was not affected by ziprasidone administration.

**Conclusions:** Behavioural abnormalities in new mothers in the postpartum period elicited as a result of STS deficiency are likely to be the culmination of many small gene expression changes. Our data are consistent with the idea that olfactory function is key to postpartum maternal behaviour in mice, and suggest that aberrant expression of olfactory system genes may partially underlie abnormal maternal behaviour in STS-deficient women.

## Introduction

Steroid sulfatase is an enzyme which cleaves sulfate groups from a variety of steroid hormones e.g. dehydroepiandrosterone sulfate (DHEAS), thereby altering their solubility and activity [Mueller et al., 2017]. STS is expressed in numerous mammalian tissues, with highest expression in the placenta [www.ncbi.nlm.nih.gov/unigene/]; in the developing and adult human brain, relatively high STS expression and activity is seen in the cortex, thalamus, cerebellum, basal ganglia, hipocampus and hypothalamus [Stergiakouli et al., 2011; Perumal and Robins, 1973]. STS deficiency is associated with increased developmental and mood disorder risk and a number of behavioural differences including: inattention, increased impulsivity and altered mood and social function [Cavenagh et al., 2019; Chatterjee et al., 2016; Trent and Davies, 2013]; these behavioural differences may be mediated, in part, by underlying changes in serotonergic or cholinergic function [Trent et al., 2013; Trent et al., 2012; Rhodes et al., 1997].

In both humans and rodents, STS expression and activity increases in brain and peripheral tissues towards the end of pregnancy and into the postpartum period; hence, enzyme deficiency or dysfunction could potentially be associated with postpartum psychopathology [Davies, 2018; Davies, 2012]. Consistent with this, we have recently shown that women who are heterozygous for genetic mutations encompassing *STS* are at increased risk of postpartum mood disorders [Cavenagh et al., 2019]. We have also demonstrated that female mice, in which STS activity is acutely, and systemically, inhibited with 667-Coumate shortly after giving birth, show altered maternal behaviour (specifically anxiety-related and startle phenotypes) relative to vehicle-treated mice; these drug-induced behavioural abnormalities can be partially reversed by concurrent administration of the atypical antipsychotic drug ziprasidone [Humby et al., 2016].

To investigate the neurobiology underlying the postpartum behaviour phenotypes in STS deficient individuals, we compared whole brain gene expression in behaviourally-defined 667-Coumate and vehicle-treated new mouse mothers. Given the large between-group behavioural differences, we suspected that screening by microarray would be able to readily identify robust gene expression differences and candidate biological pathways, and we reasoned that analysis of whole brain tissue would capture activity changes across multiple interacting brain regions.

## Methods

### Drug administration and behavioural analysis

Within 12hr of giving birth, female mice were injected with either 10mg/kg 667-Coumate or vehicle solution *per os* (n=12 per group); they were then injected with the same solution 48hr later, and behaviourally-tested 12hr after that. Behavioural testing comprised of sequential assessment on the elevated plus maze, in a locomotor activity chamber, and in a startle/prepulse inhibition paradigm. A second batch of mice had 667-Coumate injections as described above, but also ziprasidone injections i.p. 24hr after giving birth, and 1hr prior to behavioural testing at doses of either 0, 0.3 or 1.0mg/kg (n=11, n=11 and n=7 respectively). Between-group behavioural measures were compared using parametric (two-tailed unpaired t-test or One Way ANOVA) or non-parametric (Mann Whitney U test or Kruskal Wallis test) statistics depending upon normality of the data as determined by Shapiro-Wilk test. Results are presented as either mean±standard deviation of the mean or median with 95% confidence intervals determined by bootstrapping for normal and non-normal data respectively. Experiments were performed according to the UK Animal Scientific Procedures Act (1986) under Project Licence 30/3140.

### Tissue collection, RNA extraction and cDNA synthesis

3hr after behavioural testing, subjects were culled by cervical dislocation. Whole brains were immediately removed, bisected sagitally, and frozen on dry ice. High-quality total RNA was extracted from the right hemisphere of the brain using RNeasy Plus Universal Midi Kit (Qiagen) according to the manufacturer’s instructions. For microarray analysis, three biological replicates per group were generated, with each replicate containing equal amounts of RNA pooled from four hemibrains; 260/280 absorbance ratios of 2.05-2.08 and RIN numbers of 9.4-9.8 were recorded for the six replicates using Agilent 2100 Bioanalyzer (Agilent Technologies, Palo Alto, CA, USA). 20μl cDNA solution per hemibrain was synthesised from 4-5g RNA using RNA-to-cDNA EcoDry Premix with random primers (Clontech), and was diluted 50-fold with distilled water.

### Microarray hybridisation and bioinformatic analysis

Microarray analysis was conducted by Central Biotechnology Services at Cardiff University according to standard protocols. Briefly, biotin-labelled targets for the microarray experiment were prepared using 100ng of total RNA per replicate. Sense single-stranded cDNA was synthesized, fragmented and labelled using the Genechip WT PLUS Reagent Kit (Affymetrix) in conjunction with the Affymetrix Genechip Poly-A RNA Control Kit as described in the User Manual (P/N 703174). A hybridization cocktail containing the biotinylated target was incubated with the GeneChip Mouse 2.0 ST array (Affymetrix) at 60rpm for 16hr at 45°C in a Genechip Hybridisation Oven 645. After hybridization, non-specifically bound material was removed by washing, and specifically-bound target was detected using the GeneChip Hybridization, Wash and Stain Kit, in conjunction with the GeneChip Fluidics Station 450 (Affymetrix). The arrays were scanned using a GeneChip Scanner 3000 7G (Affymetrix) in conjunction with Affymetrix Genechip Command Console (AGCC) software. Data were normalised using Robust Multiarray Average (RMA); the overall pattern of normalised expression data appeared equivalent across the six replicates. Differential gene expression analysis was performed using lmFit and eBayes from the Limma package [Sartor et al., 2006]. FDR-corrected and nominal p-values were generated for each probe, with values<0.05 being regarded as nominally statistically-significant. Pathway analysis was performed using Database for Annotation, Visualisation and Integrated Discovery version 6.8 (DAVID, RRID: SCR_001881) [Dennis et al., 2003]. Raw microarray data are available in the ArrayExpress database (http://www.ebi.ac.uk/arrayexpress, RRID: SCR_002964) under accession number E-MTAB-7233.

### Quantitative Polymerase Chain Reaction (qPCR)

qPCR was performed using a Rotorgene 6000 coupled with a CAS1200 automated set-up, and utilizing standard consumables (Qiagen, Manchester, UK). PCR reactions were performed using 5μl cDNA mix and 200nM custom-designed primers (Table 1) and SensiMix (Bioline, London, UK). qPCR data were analysed using ΔC_t_ methods as described previously [Isles et al., 2004] with normalisation to the mean of three ‘housekeeping gene standards’ (*Hprt*, *Gapdh* and *Rn18s*) whose expression was significantly (p<0.001) correlated within the samples. Groups were compared by either Mann-Whitney U test, unpaired t-test or One Way ANOVA.

**Table 1.**
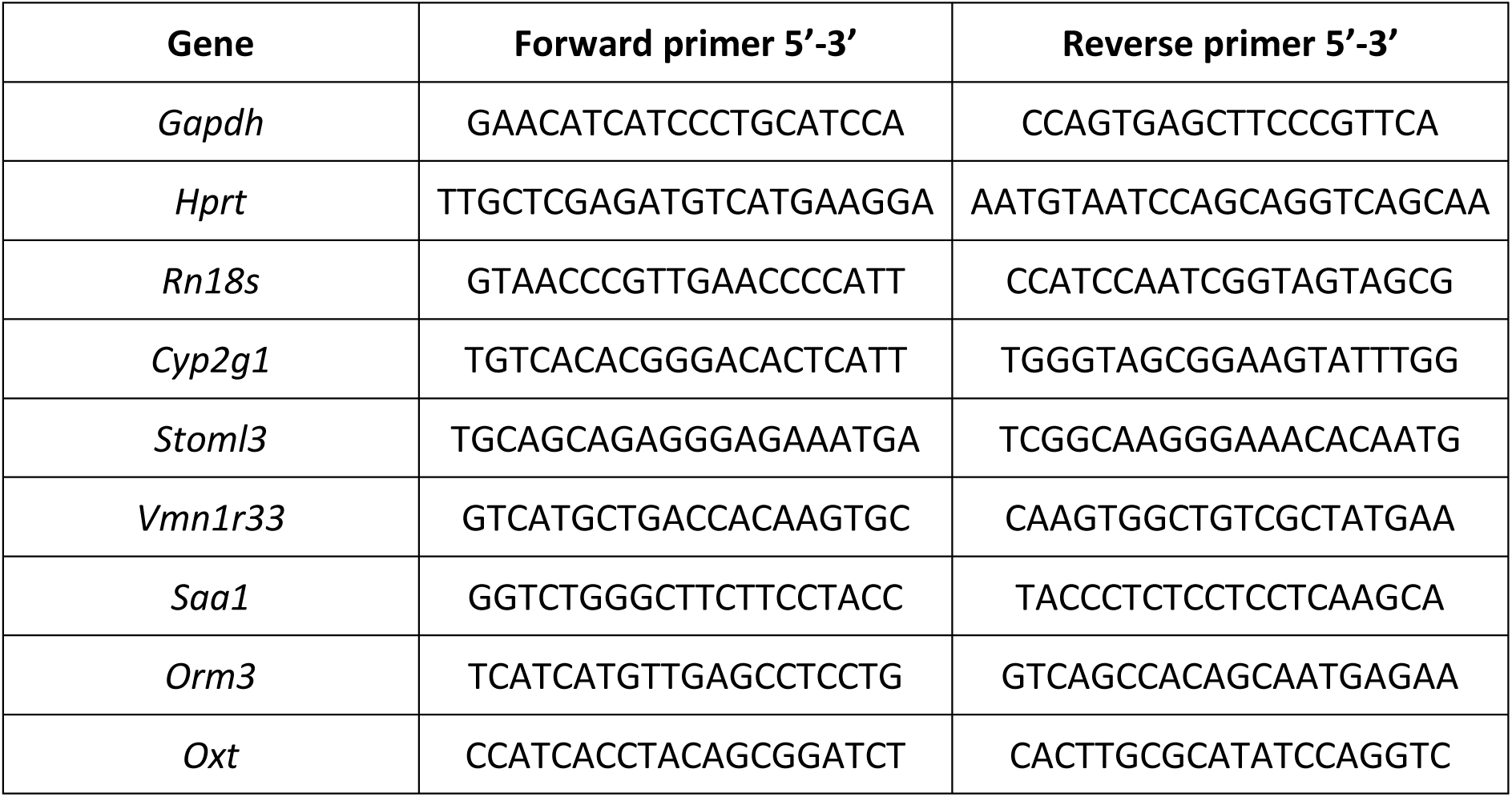
Primer sequences used for quantitative PCR analysis. Primers were designed to allow optimum amplification, to span intron-exon boundaries where possible, and to amplify key coding gene transcripts.

## Results

### Behavioural and endocrine comparison in animals used for microarray analysis

A subset of behaviourally-defined animals from Humby et al. (2016) were selected for gene expression analysis. 667-Coumate and vehicle-treated mothers differed significantly with respect to numbers of rears on the elevated plus maze (57 (95%CI:19-86.5) vs. 0 (95%CI:0-0), Mann-Whitney U=10.5, p<0.0001), latency to enter the open arm of the elevated plus maze (2.2(95%CI:2.0-3.0)s vs. 8.5s(95%CI:3.2-12.2), Mann-Whitney U=28.0, p=0.01), and startle response at 120dB covarying for bodyweight and baseline activity (51.6±8.7 vs. 62.5±8.3 abitrary units, t[1,20]=4.61, p=0.04). These group differences were not confounded by differential locomotor activity as indexed by infra-red beam breaks per hour (1118 (95%CI:925-2284) vs. 1600 (95%CI:786-2143), Mann-Whitney U=72.0, p=1.0), nor by number of live pups born to each mother (7 (95%CI:6-8) vs. 7(95%CI:6-8), Mann-Whitney U=71.0, p=0.95). The median serum DHEAS:DHEA ratio in 667-Coumate-treated mothers (n=11) was ∼1.6-fold higher than that in vehicle-treated mothers (n=5)(333 vs. 211), indicating efficacy of systemic STS inhibition by the drug.

### Significantly-differentially expressed genes in the microarray analysis

We identified 1,081 unique differentially-expressed transcripts with a nominal p-value cut-off<0.05; the 24 most highly differentially expressed of these (fold change>1.5) are listed in Table 2. No significantly differentially expressed genes were identified using the comparatively stringent cut-off of False Discovery Rate (FDR)-corrected p-value of <0.1.

**Table 2.**
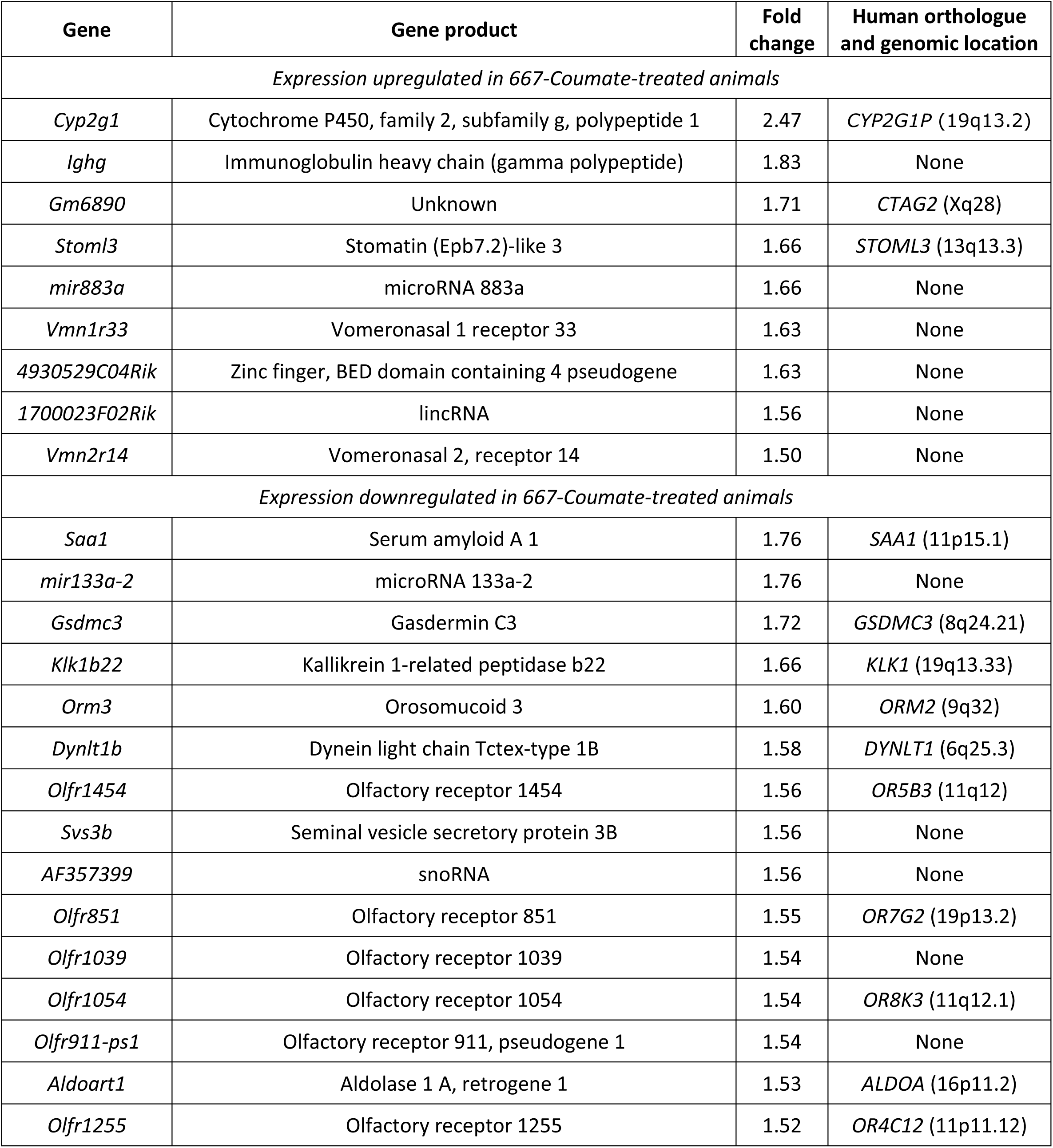
Most highly-differentially expressed genes (>1.5 fold change, p<0.05) between vehicle and 667-Coumate-treated whole mouse brain according to microarray analysis

### Pathway analysis

Pathway analysis of the 1,081 differentially-expressed genes at p<0.05 with the mouse genome as background identified only one KEGG (Kyoto Encyclopedia of Genes and Genomes [Ogata et al., 1999]) pathway, olfactory transduction, as being significant (Benjamini-corrected p=1.8×10^−3^). The following gene ontology (GO) terms for biological pathway were significant after correction for multiple testing (G-protein coupled receptor signalling p=1.2×10^−3^ and sensory perception of smell p=7.5×10^−4^), as were the following GO terms for molecular function (G-protein coupled receptor activity p=5.1×10^−5^ and olfactory receptor activity p=1.3×10^−4^). We tested the possibility that identification of the olfactory pathway could have been artefactual due to the relatively low and variable expression of the associated genes, by filtering for low expression variability within the following R Package using the nsFilter function with default value of var.cutoff 0.5: https://www.rdocumentation.org/packages/genefilter/versions/1.54.2/topics/nsFilter. This analysis resulted in just 213 unique genes that were differentially-expressed at a nominal p-value<0.05, 35 of which had a fold change in expression >1.5 (the 24 genes from Table 2 as well as *Psg25*, *1700047G07Rik*, *Ugt2b5*, *Vmn1r86*, *Olfr493*, *Olfr295*, *Vmn1r37*, *Olfr800* and *Fpr2*); for the 213 genes, the KEGG pathway ‘olfactory transduction’ remained highly significant (p=3.6×10^−11^), as did the GO biological pathway terms ‘G-protein-coupled receptor signalling’ and ‘sensory perception of smell’ (p=7.9×10^−12^ and p=1.2×10^−12^ respectively) and the GO molecular function terms ‘G-protein coupled receptor activity’ and ‘olfactory receptor activity’ (p=9.6×10^−17^ and 2.4×10^−13^ respectively).

### Quantitative PCR (qPCR)

qPCR analysis was performed for five of the most highly-differentially expressed genes from Table 2, as well as for *Oxt* (downregulated 1.33-fold, nominal p<0.005 in the microarray analysis). *Oxt* encodes the oxytocin protein important in maternal physiology, the abnormal expression of which has been implicated in postpartum mood disorders [Moura et al., 2016; Gammie et al., 2016]. Whilst the direction of expression difference was generally consistent across the microarray and qPCR analyses for these genes, only two genes (*Cyp2g1* and *Stoml3*) were significantly differentially-expressed according to qPCR (U=34.0, one-tailed p=0.014 and t[15.3]=-1.83, one-tailed p=0.04 respectively)(Figure 1); for the remaining four genes, p>0.075. We next tested whether *Cyp2g1* and *Stoml3* expression was sensitive to the co-administration of vehicle or ziprasidone (0.3mg/kg or 1.0mg/kg) in 667-Coumate treated animals; the expression of neither of these genes was affected: *Cyp2g1* (F[2,28]=0.58, p=0.57) and *Stoml3* (F[2,28]=2.83, p=0.08)(Figure 2).

**Figure 1.**
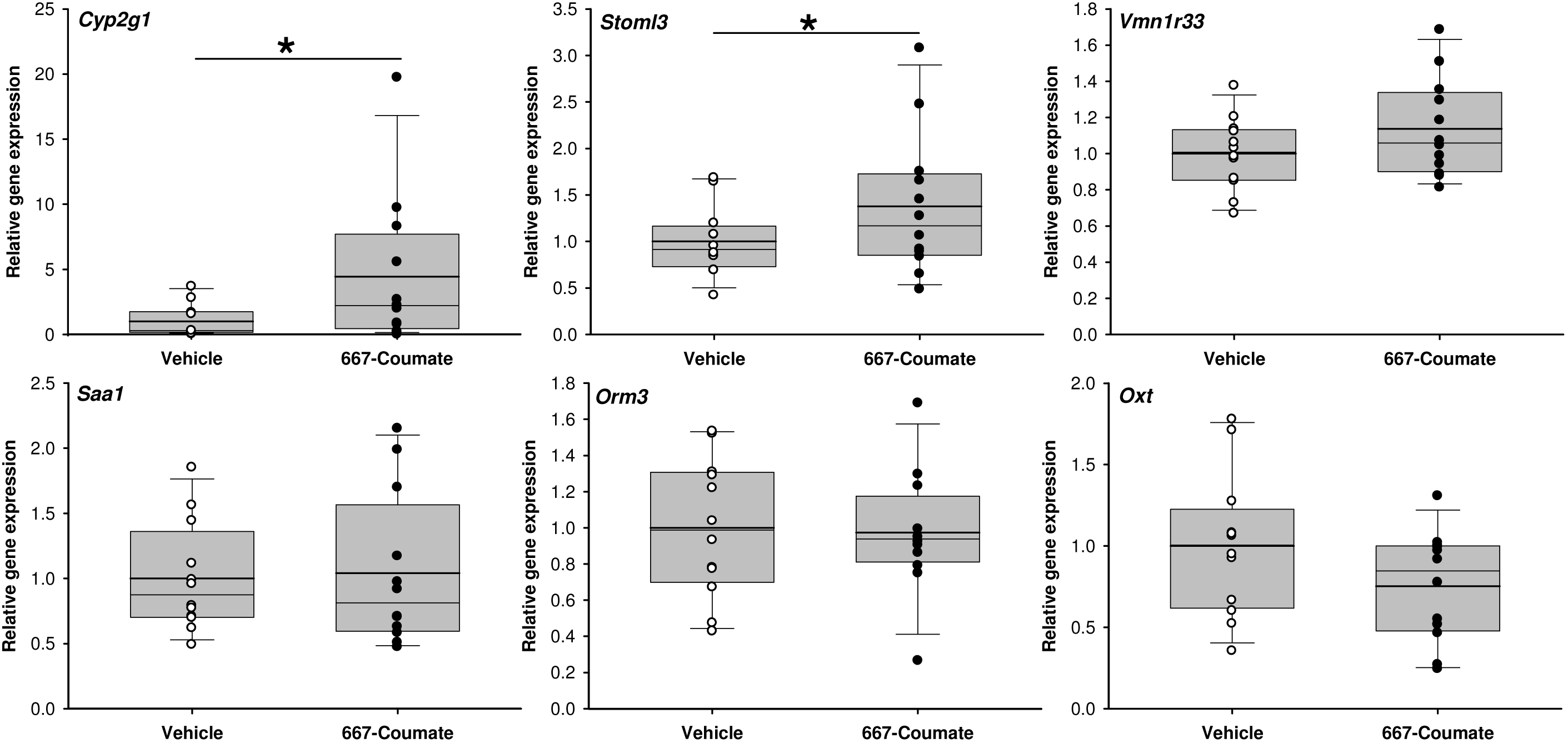
Comparison of gene expression in vehicle and 667-Coumate treated whole mouse brain by quantitative PCR. Boxes show interquartile range with median and mean (bold line) values, and whiskers represent 5% and 95% confidence intervals. *p<0.05.

**Figure 2.**
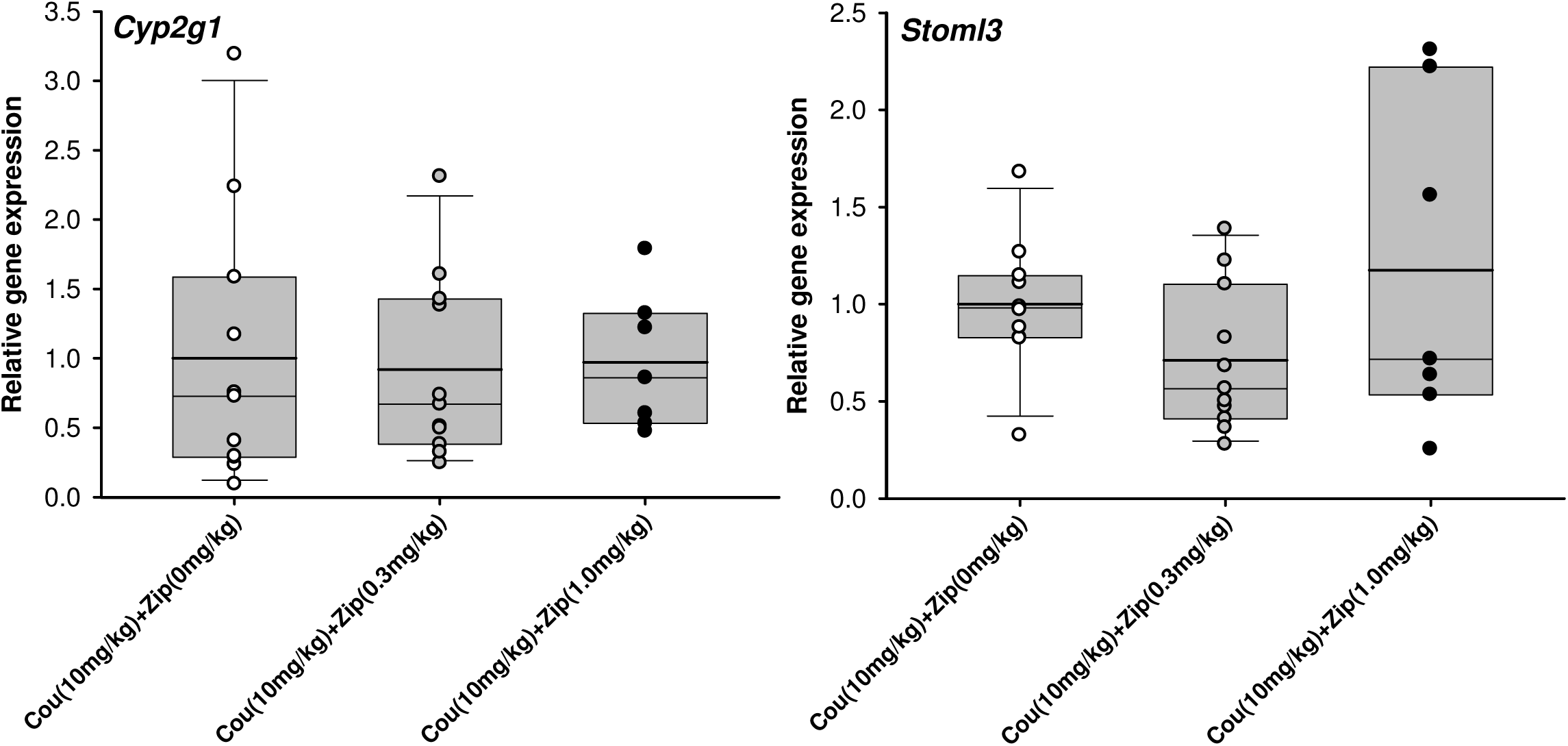
Comparison of gene expression in whole mouse brain by quantitative PCR in animals administered 667-Coumate and either 0, 0.3 or 1.0mg/kg ziprasidone. Boxes show interquartile range with median and mean (bold line) values, and whiskers represent 5% and 95% confidence intervals.

## Discussion

In this study we investigated whole brain gene expression signatures underlying abnormal maternal behaviour in a novel mouse model of postpartum mood disorder (STS inhibition with 667-Coumate). Although our vehicle and 667-Coumate treated groups differed substantially both in terms of their behaviour and with respect to a peripheral endocrine marker of STS inhibition (DHEAS:DHEA ratio), and the microarray experiment was performed with standard quality control procedures, we identified surprisingly few gene expression differences between the groups; those changes which we did identify by microarray were relatively small in magnitude and most could not be verified by quantitative PCR. These data indicate that, at least at the timepoint we assayed, large brain gene expression differences do not substantially contribute towards abnormal maternal behaviour in the STS inhibition mouse model, and suggest that the behavioural differences are associated with another underlying biological mechanism e.g. the aggregate effect of small expression changes across many genes. This idea is consistent with our previous observations of: a) few large, statistically-significant, gene expression differences between whole brain samples from male mice lacking the *Sts* gene and wildtype animals [Trent et al., 2014], despite considerable between-group behavioural differences [Davies et al., 2014; Trent et al., 2013; Trent and Davies, 2013; Trent et al., 2012; Davies et al., 2009] and b) evidence that small brain expression changes (<1.5-fold) detectable between 667-Coumate and vehicle-treated mice by qPCR, but not detectable by the present microarray study, might be associated with postpartum behavioural phenotypes [Humby et al., 2016]. We did not identify any overlap between genes significantly differentially expressed in the current study, and those whose expression was altered in whole brain from *Sts*-deficient male mice [Trent et al., 2014], and nor was there any overlap with genes whose skin expression was altered in male patients with steroid sulfatase deficiency [Hoppe et al., 2012]; hence, the genetic mechanisms associated with acute STS inhibition in females and constitutive STS deficiency in males are likely to be largely dissociable.

We did observe robust upregulation of the *Cyp2g1* and *Stoml3* genes (∼3-fold and 1.5-fold respectively) in 667-Coumate treated whole brain. The expression of both genes is restricted to the olfactory system in mice [Zhuo et al., 2004; Goldstein et al., 2003]. CYP2G1 is a major P450 enzyme in the olfactory mucosa of rodents, and, whilst its absence in homozygous knockout mice does not appear to impair olfaction, it does result in altered steroid hormone metabolism and metabolic activation of coumarin [Zhuo et al., 2004]. Increased expression of *Cyp2g1* following administration of 667-Coumate, a tricyclic coumarin sulfamate [Woo et al., 2000], is perhaps, therefore, unsurprising, and further evidence that the drug is influencing neurophysiology. In man, functional *Cyp2g1* orthologues appear to be rare [Sheng et al., 2000]. Whilst the biological roles of STOML3 remain to be fully clarified, there is some evidence that the protein mediates mechanosensory processes [Wetzel et al., 2007]. The elevated expression of *Cyp2g1* and *Stoml3* genes in 667-Coumate-treated mouse brain could not be reduced through antipsychotic (ziprasidone) adminstration, indicating that these genes and their asociated proteins are unlikely to play a large role in mediating the rescue effect of ziprasidone on aspects of maternal behaviour.

Pathway analysis incorporating all nominally-significant microarray hits (including those with low differential expression) suggested that 667-Coumate may perturb olfactory transduction processes and that this perturbation may mediate drug-induced effects on postpartum maternal behaviour in the mouse; potentially, the increased prevalence of postpartum mood disorder in STS deficient women may be partially attributable to abnormalities within the olfactory system and/or its links to the limbic system. This possibility is feasible in light of the clinical observation that 667-Coumate administration can elicit taste disturbances in female patients [Stanway et al., 2006], and is consistent with an extensive literature on the importance of olfactory (and associated limbic) function in mammalian mothers [Corona and Levy, 2015], with evidence that several steroid sulfates act as ligands within the mouse accessory olfactory system [Meeks et al., 2010], with the expression of multiple olfactory receptors within human brain tissue [Flegel et al., 2013], and with recent findings that olfactory processes are perturbed in a genetic mouse model of abnormal maternal behaviour [Creeth et al., 2018] and multiple mood disorders [Kamath et al., 2018]. With regard to specific candidate genes of interest, both the study by Creeth *et al*. and the present study highlighted *Olfr59* expression as being significantly upregulated (∼1.4-fold, p<0.05) in hypothalamus and whole brain respectively in mothers with maternal behavioural abnormalities; this gene has no clear human orthologue.

The present study is limited in two main ways. First, we examined gene expression across the whole brain. Whilst this strategy is useful for capturing widespread expression changes and is sensible given our lack of knowledge about the underlying neuroanatomy of postpartum mood disorders, it is unlikely to detect gene expression changes between groups that are regionally-specific. Second, microarray technology, whilst cheap and easily implementable, is relatively insensitive to small gene expression changes, and has low resolution with regard to determining differentially expressed gene transcripts [Zhao et al., 2014]. Hence, future between-group comparisons might use a more specific, and more sensitive, technique such as RNA sequencing to assay gene expression in selected brain regions of importance to maternal behaviour and mood regulation, and whose chemistry is known to be altered by STS deficiency e.g. hippocampus.

## Acknowledgements

We are grateful to staff at the Central Biotechnology Services (CBS) and Core Bioinformatics and Statistics Team within the College of Biomedical and Life Sciences at Cardiff University for their help with performing and analysing the microarray experiments. The work was performed within the Medical Research Council Centre for Neuropsychiatric Genetics and Genomics (Centre grant MR/L010305/1) and was funded by a Neuroscience and Mental Health Research Institute, Cardiff University seedcorn grant to TH and WD. This manuscript has been released as a preprint at bioRxiv [Humby and Davies, 2019].

